# Platon: identification and characterization of bacterial plasmid contigs in short-read draft assemblies exploiting protein-sequence-based replicon distribution scores

**DOI:** 10.1101/2020.04.21.053082

**Authors:** Oliver Schwengers, Patrick Barth, Linda Falgenhauer, Torsten Hain, Trinad Chakraborty, Alexander Goesmann

## Abstract

Plasmids are extrachromosomal genetic elements replicating independently of the chromosome which play a vital role in the environmental adaptation of bacteria. Due to potential mobilization or conjugation capabilities, plasmids are important genetic vehicles for antimicrobial resistance genes and virulence factors with huge and increasing clinical implications. They are therefore subject to large genomic studies within the scientific community worldwide. As a result of rapidly improving next generation sequencing methods, the amount of sequenced bacterial genomes is constantly increasing, in turn raising the need for specialized tools to (i) extract plasmid sequences from draft assemblies, (ii) derive their origin and distribution, and (iii) further investigate their genetic repertoire. Recently, several bioinformatic methods and tools have emerged to tackle this issue; however, a combination of both high sensitivity and specificity in plasmid sequence identification is rarely achieved in a taxon-independent manner. In addition, many software tools are not appropriate for large high-throughput analyses or cannot be included into existing software pipelines due to their technical design or software implementation. In this study, we investigated differences in the replicon distributions of protein-coding genes on a large scale as a new approach to distinguish plasmid-borne from chromosome-borne contigs. We defined and computed statistical discrimination thresholds for a new metric: the replicon distribution score (RDS) which achieved an accuracy of 96.6%. The final performance was further improved by the combination of the RDS metric with heuristics exploiting several plasmid specific higher-level contig characterizations. We implemented this workflow in a new high-throughput taxon-independent bioinformatics software tool called Platon for the recruitment and characterization of plasmid-borne contigs from short-read draft assemblies. Compared to PlasFlow, Platon achieved a higher accuracy (97.5%) and more balanced predictions (F1=82.6%) tested on a broad range of bacterial taxa and better or equal performance against the targeted tools PlasmidFinder and PlaScope on sequenced *E. coli* isolates. Platon is available at: platon.computational.bio

**Data Summary:**

1. Platon was developed as a Python 3 command line application for Linux.
2. The complete source code and documentation is available on GitHub under a GPL3 license: https://github.com/oschwengers/platon and platon.computational.bio.
3. All database versions are hosted at Zenodo: DOI 10.5281/zenodo.3349651.
4. Platon is available via bioconda package `platon`
5. Platon is available via PyPI package `cb-platon`
6. Bacterial representative sequences for UniProt’s UniRef90 protein clusters, complete bacterial genome sequences from the NCBI RefSeq database, complete plasmid sequences from the NCBI genomes plasmid section, created artificial contigs, RDS threshold metrics and raw protein replicon hit counts used to create and evaluate the marker protein sequence database are hosted at Zenodo: DOI 10.5281/zenodo.3759169
7. 24 *Escherichia coli* isolates sequenced with short read (Illumina MiSeq) and long read sequencing technologies (Oxford Nanopore Technology GridION platform) used for real data benchmarks are available under the following NCBI BioProjects: PRJNA505407, PRJNA387731

**Impact Statement:** Plasmids play a vital role in the spread of antibiotic resistance and pathogenicity genes. The increasing numbers of clinical outbreaks involving resistant pathogens worldwide pushed the scientific community to increase their efforts to comprehensively investigate bacterial genomes. Due to the maturation of next-generation sequencing technologies, nowadays entire bacterial genomes including plasmids are sequenced in huge scale. To analyze draft assemblies, a mandatory first step is to separate plasmid from chromosome contigs. Recently, many bioinformatic tools have emerged to tackle this issue. Unfortunately, several tools are implemented only as interactive or web-based tools disabling them for necessary high-throughput analysis of large data sets. Other tools providing such a high-throughput implementation however often come with certain drawbacks, *e*.*g*. providing taxon-specific databases only, not providing actionable, *i*.*e*. true binary classification or achieving biased classification performances towards either sensitivity or specificity.

Here, we introduce the tool Platon implementing a new replicon distribution-based approach combined with higher-level contig characterizations to address the aforementioned issues. In addition to the plasmid detection within draft assemblies, Platon provides the user with valuable information on certain higher-level contig characterizations. We show that Platon provides a balanced classification performance as well as a scalable implementation for high-throughput analyses. We therefore consider Platon to be a powerful, species-independent and flexible tool to scan large amounts of bacterial whole-genome sequencing data for their plasmid content.

## Introduction

Plasmids are bacterial extrachromosomal DNA elements, which replicate independently of the chromosome. They are mostly circular, have characteristic copy numbers per cell and carry genes that are usually not essential under normal conditions but rather allow bacteria to adapt to specific environments and conditions [1]. These genes, for instance, provide antibiotic or heavy metal resistances, are involved in alternative metabolic pathways or encode for virulence factors [2]. As plasmids are not only inherited by daughter cells, but can also be dispersed by horizontal gene transfer, they can spread rapidly within and between bacterial populations [3–5]. For example, identical antibiotic resistance plasmids have already been isolated from humans and animals [6]. Thus, plasmids are important mediators of antibiotic resistance spread and recent findings confirm they frequently play a major role in clinical outbreaks [7, 8]. Therefore, it is of huge importance to properly identify and analyze plasmids.

Such analysis can be performed by plasmid DNA isolation followed by sequencing [9]. However, due to decreased sequencing costs, nowadays it is affordable and often easier to sequence the entire genome of bacterial organisms by next-generation whole-genome shotgun sequencing [10]. Furthermore, this approach allows the re-analysis of already sequenced genomes to identify plasmids which have not been detected before. Unfortunately, this introduces a new issue which needs to be addressed: Plasmid and chromosomal contigs are both mixed in draft assemblies and need to be distinguished from each other.

This task, however, is hard to achieve due to biological and technical reasons [11]. Plasmids often contain mobile genetic elements, *e*.*g*. transposons and integrons, which are drivers for the genetic exchange between different DNA replicons and regions [12, 13]. Hence, these genetic elements are often encoded on both replicon types and thus, the origin of DNA fragments encoding such elements is hard to predict. Modern short-read assemblers pose an additional intricacy further aggravating these issues as they are notoriously hard pressed to correctly assemble repetitive regions such as aforementioned DNA elements [14]. To tackle this issue, many new bioinformatic tools have recently been developed, following different approaches: (i) Recycler and plasmidSPAdes [15, 16] exploit coverage variations of sequenced DNA fragments within a genome; (ii) PLACNET investigates paired-end reads linking contig ends [17]; (iii) PlasmidFinder searches for plasmid specific motifs, *i*.*e*. incompatibility groups [18]; (iv) cBar, PlasFlow and mlPlasmids use machine learning methods to classify k-mer frequencies [19–21]; (v) PlaScope and PlasmidSeeker perform fast k-mer-based database searches of known plasmid sequences [22, 23]; Recycler additionally exploits information on circularization [15]. Overall, each approach provides unique advantages and drawbacks. For example, approaches based on sequencing coverage variations are unable to detect plasmids with copy numbers equal to the chromosome whereas sequence motif and k-mer-based methods tend to identify only known plasmids. This often leads to distinct profiles in terms of sensitivity and specificity which are often biased towards one of both metrics and as this impacts on the conducted analysis a choice must be made between conservative or more aggressive classifications [11].

A further aspect of growing importance is Big Data awareness. Due to increasing amounts of generated sequence data [24], there is a rising need for automated high-throughput analysis tools. Unfortunately, not all currently available bioinformatics software tools are suitable for high-throughput analysis, let alone the technical integration into larger analysis pipelines [25–27] due to interactive designs or web-based implementations [17, 18, 21, 28]. Taxon-specific database designs also pose additional barriers as users might not have sufficient computational resources or bioinformatics support to build customized or large multi-taxon databases [20, 22]. Furthermore, dependence on raw or intermediate data such as sequence reads and assembly graphs might impede analyses as such data might not be available [15, 16]. In order to allow for Big Data scaling necessities, bioinformatic software tools should therefore be designed and implemented in a high-throughput savvy manner, including: (i) where possible a taxon-independent database design; (ii) a non-interactive command line implementation; (iii) an actionable classification output, *i*.*e*. a true binary classification.

To address these issues we present Platon, a new bioinformatics software tool to distinguish and characterize plasmid contigs from chromosome contigs in bacterial draft assemblies following a new approach: analysis of replicon distribution differences of protein-coding genes, *i*.*e*. frequency differences of being encoded on plasmid or chromosome contigs. The rationale behind this protein sequence-based replicon, *i*.*e*. chromosome *vs*. plasmid, classification is a natural distribution bias of certain protein-coding genes. For instance, essential housekeeping genes mandatory for bacterial organisms are mostly encoded on chromosomes [2]. In contrast, genes providing an evolutionary advantage under distinct situations are rather widespread on plasmids, *e*.*g*. antibiotic resistance and virulence genes. Here, we introduce “replicon distribution scores” (**RDS**), a new metric to express the measured bias of protein-coding genes’ replicon distributions to distinguish plasmid from chromosome related contigs.

## Methods and Implementation

### Marker protein sequences and computation of replicon distribution scores

To build a database of marker protein sequences (**MPS**) we collected all bacterial representative sequences of UniProt’s UniRef90 protein clusters (n=69,803,841) [29] and analyzed their replicon distributions, *i*.*e*. the normalized plasmid vs. chromosome abundance ratios. Therefore, we conducted a homology search via Diamond [30] of all MPS against coding sequences (CDS) predicted via Prodigal [29] on two reference replicon sets, *i*.*e*. all NCBI plasmid sequences from the bacterial NCBI Genomes plasmid section (n=17,369) (ftp://ftp.ncbi.nlm.nih.gov/genomes/GENOME_REPORTS/plasmids.txt) and chromosomes of all complete bacterial NCBI RefSeq release 98 genomes. To prevent potential plasmid contamination in the chromosome set, all replicons shorter than 100 kbp were excluded resulting in 17,430 chromosome sequences. Resulting alignment hit counts (A) of the single best hit per sequence with a sequence coverage larger than or equal to 80% and a sequence identity of at least 90% as well as the number of replicons (R) for both plasmids (p) and chromosomes (c) were integrated into a normalized, transformed and scaled replicon distribution score RDS for each cluster defined by:

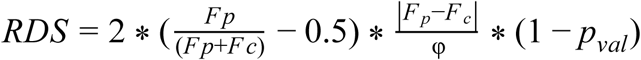

with 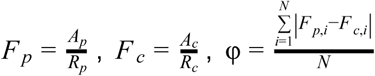, N the number of elements of the MPS database and p_val_ the p value of a two-sided Fisher’s exact test using a contingency table of hit and no-hit counts for both replicon types.

Thus, the RDS value of a protein sequence represents its replicon distribution bias as both ratio and absolute difference of hit count frequencies as well as its statistical power, *i*.*e*. 1 - p value. As a first factor of the formula, the hit count frequency ratio 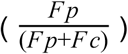 is transformed by the minuend −0.5 and the factor 2 to the range [-1,1] and hence, shifts RDS values of chromosomal proteins to a negative range [-1,0] and to a positive value range [0,1] for proteins with a positive plasmid bias. To integrate the scale of the difference in the hit count frequencies we added the absolute difference of replicon hit count frequencies (*F*_*p*_ − *F*_*c*_) normalized to the mean difference of hit count frequencies of all MPS (φ) as a second factor. In order to also include a measure of statistical confidence into the new RDS metric a third factor (1 − *p*_*val*_) was added, taking the p value of a two-sided Fisher’s exact test using a contingency table of hit and non-hit counts of both replicon types under the assumption that these are not equally distributed - the main idea behind the RDS metric. Thus, RDS values resulting from statistically insignificant hit count numbers get minimized towards zero. In order to finally classify entire contigs, the mean RDS of all per-protein-sequence RDS values of each contig is calculated and then tested against defined thresholds. Predicted CDS, for which no MPS can be identified get assigned the neutral default RDS value of zero.

### Evaluation of replicon distribution scores

In order to assess the discriminative power of protein sequence based RDS, we created 10 random fragments of each sequence in the reference replicon sets for each of the following lengths: 1 kbp, 5 kbp, 10 kbp, 20 kbp and 50 kbp. For each random fragment, a mean RDS was computed and tested against a range of discrimination thresholds between −50 and 10 with a 0.1 step size. For each discrimination threshold a confusion matrix was set up upon which sensitivity (*tp/(tp*+*fn)*), specificity (*tn/(tn*+*fp)*) and accuracy (*(tp*+*tn)/(tp*+*tn*+*fp*+*fn)*) metrics were calculated, where `tp`, `tn`, `fp` and `fn` is the number of true positives, true negatives, false positives and false negatives, respectively.

### Higher-level contig characterization

The comprehensive characterization of contigs by higher-level plasmid-related sequence analysis often requires many specialized command line and web-based tools and thus is a time-consuming task. To streamline this process, we implemented and included many higher-level sequence analyses in the workflow. Hence, Platon provides valuable contig information and can take advantage thereof by integrating applied heuristics into the classification process. Contigs are comprehensively characterized comprising different approaches: (i) test for circularization; (ii) detection of incompatibility groups; (iii) detection of rRNA genes; (iv) detection of antimicrobial resistance genes; (v) homology search against reference plasmid sequences; (vi) detection of oriT sequences; (vii) detection of plasmid replication genes; (viii) detection of mobilization genes; (ix) detection of conjugation genes.

Contigs are tested for circularization by aligning subsequences from both ends against each other using nucmer from the MUMmer package [31]. Contig ends with overlaps larger than or equal to 100 bp and an identity larger than 95% are considered circularizable. To detect incompatibility groups, Platon conducts a homology search using the PlasmidFinder database (n=273) [18] via BLAST+ [32] against contigs filtering for query coverages larger than or equal to 60% and percent sequence identities larger than 90%. Although there are rare exceptional cases described in the literature [33], the majority of ribosomal genes are encoded on chromosomes [33]. In order to exploit this distribution bias, ribosomal genes are detected via Rfam and Infernal [34]. As antimicrobial resistance genes are often encoded on mobile genetic elements as for instance plasmids, Platon uses the NCBI ResFam hidden markov models (**HMM**) database [35] and HMMER [36] to detect potential antimicrobial resistance genes. In order to detect contigs as subsequences of larger plasmids or entire plasmids with known sequences, Platon conducts a homology search via BLAST+ [30] against the RefSeq plasmid sequence database [37] filtering for query coverages and percent sequence identities larger than or equal to 80% setting a dynamic `-word_size` parameter to 1% of the query contig length. To detect oriT sequences, Platon conducts a BLAST+ [32] homology search against oriT sequences of the MOB-suite [38] database filtering for both 90% sequence coverage and identity.

Depending on its genetic backbone, plasmids can be mobilizable or conjugative [4]. The presence or absence of specialized proteins involved in the replication, mobilization and conjugation processes play important roles as determinants for the classification of plasmids. Platon takes advantage of the highly plasmid-specific nature of these proteins by scanning predicted CDS against a custom HMM database. Therefore, we extracted relevant RefSeq PCLA protein clusters via text-mining and subsequently built HMM models on aligned protein sequences per cluster (**S1 Table**), creating two distinct HMM databases: replication and conjugation comprising 257 and 1,663 HMM models, respectively. To take advantage of the expert knowledge and manual efforts which led to the high-quality relaxase HMM profiles of the MOBscan [39] database, these were incorporated into this workflow. A scan against each HMM database is integrated into the classification process.

### Platon analysis workflow

Platon combines the analysis of the replicon distribution bias of protein sequences with a set of higher-level contig characterizations to predict the replicon origin of contigs (**Fig 1**). In a first step, Platon classifies all contigs with a length smaller than 1 kbp or larger than 500 kbp as chromosomal. The rationale behind this heuristic is that sequences with less than 1 kbp seldom host either a CDS or other exploitable information which would permit reliable classifications. On the other hand, from our experience contigs larger than 500 kbp rarely or never originate from plasmids as those often encode genetic features hindering the assembly of larger sequences such as for example transposons and integrons. Thus, this heuristic enhances the overall analysis runtime performance without unduly sacrificing classification performance.

In a second step, CDS are predicted via Prodigal [40] and searched against a database of MPS via Diamond [30] applying rigorous detection cutoffs in line with the cluster specifications of the underlying UniRef90 clusters, *i*.*e*. a coverage of at least 80% and a sequence identity of at least 90%. For each contig, the mean RDS of all detected MPS is computed. Contigs with a mean RDS lower than a sensitivity threshold (**SNT**) are classified as chromosomal sequences. Remaining contigs are then comprehensively characterized as described in the former section.

**Fig. 1.**
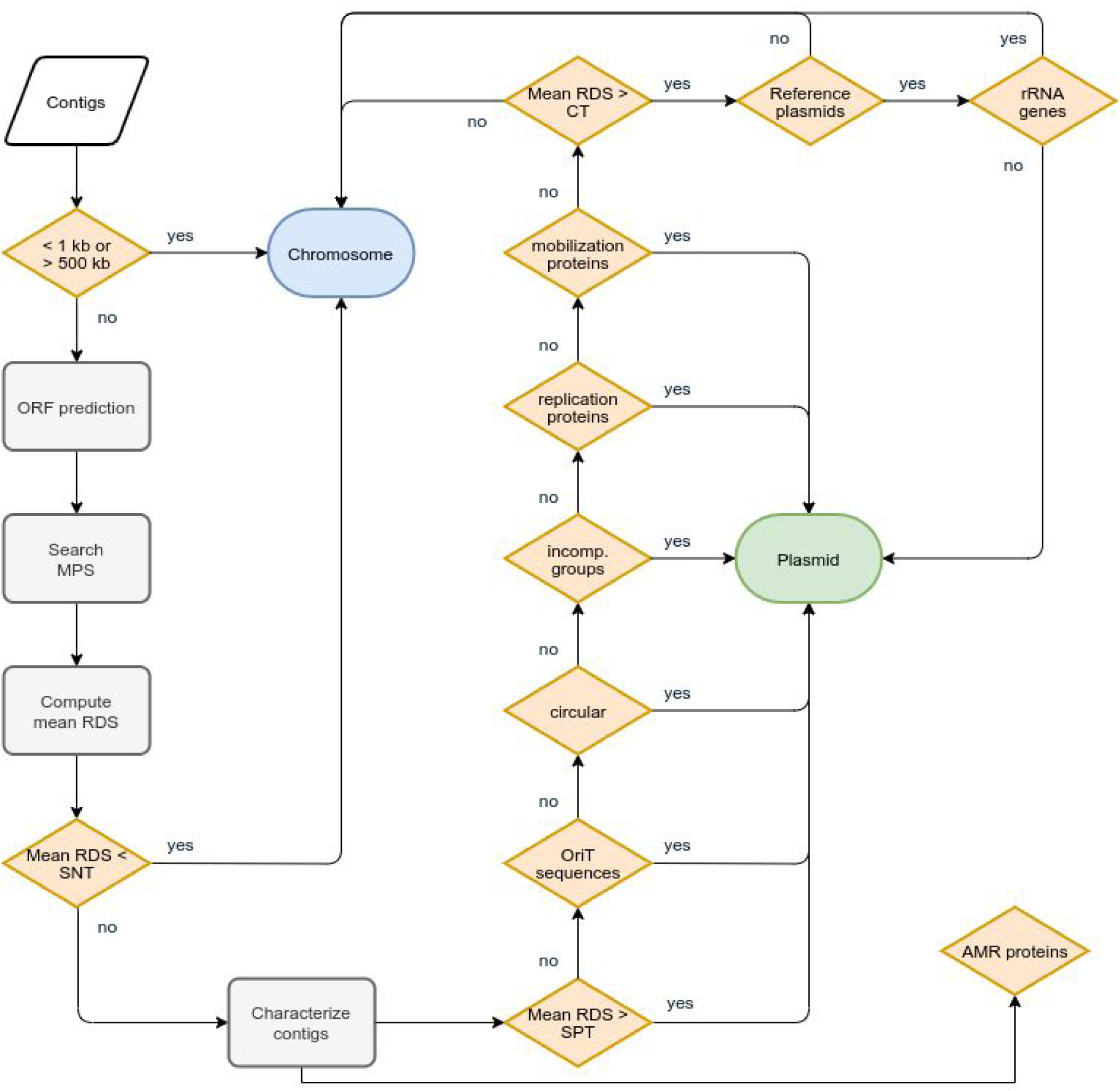
Flowchart describing the workflow implemented in Platon. ORF: open reading frames; MPS: marker protein sequences; RDS: replicon distribution score; SNT: sensitivity threshold; SPT: specificity threshold; incomp. groups: incompatibility groups; CT: conservative threshold.

Contigs are subsequently classified as plasmid sequences if one or more of the following conditions are met: the contig (i) has a mean RDS larger than a specificity threshold (**SPT**); (ii) can be circularized; (iii) provides at least one replication or mobilization protein; (iv) contains an incompatibility factor; (v) contains an oriT sequence; (vi) has a mean RDS larger than a conservative threshold (**CT**) and a BLAST+ [32] hit against the RefSeq plasmid database without encoding ribosomal genes.

### Performance benchmarks

The overall replicon classification performance of Platon v1.3.1 was benchmarked against PlaScope 1.3.1, PlasFlow 1.1.0 and the PlasmidFinder database (version 2018-11-20) in two setups: a targeted benchmark comparing Platon against PlaScope and PlasmidFinder on sequenced *Escherichia coli* isolates and an untargeted benchmark comparing Platon against PlasFlow on simulated short-read assemblies of all complete RefSeq genomes. PlaScope and PlasFlow were used with default parameters and publicly provided pre-built databases. As PlasmidFinder is currently only available as a web tool or via Docker which is not eligible on our HPC cluster setup, its workflow was reimplemented in bash using equal BLAST+ parameters (`-perc_identity 90`; query coverage>=60%). As both PlaScope and PlasFlow allow a third classification label, *i*.*e*. `unclassified` and thus, are not true binary classifiers, replicon fragments were treated as being classified as chromosomes as long as they were not explicitly classified as plasmid for the sake of comparability. For each benchmark, we calculated sensitivity, specificity and accuracy metrics as described above. To also include statistically balanced metrics, we calculated the positive predictive power (**PPV**) (*tp/(tp*+*fp)*), the negative predictive power (**NPP**) (*tn/(tn*+*fn)*) as well as F1 score and Matthews correlation coefficient (**MCC**) using the SciKit-learn Python package. For the simulated benchmark dataset, we used all bacterial NCBI RefSeq genomes (release 98) at the assembly level *Complete Genome* (n=13,930) to generate artificial short reads via ART (2.5.8) [41] with read lengths of 150 bp, 40 fold coverage, 500 bp mean insert size and 10 bp insert size standard deviation. Simulated reads were then assembled with SPAdes (3.12.0) [42] using the *--careful* and *--cov-cutoff auto* parameters. Resulting contigs (n=820,932) were aligned against original genomes with BLAST+ (2.7.1) [32] and finally labeled either as chromosome or plasmid according to the single best BLAST+ hit.

To benchmark on real data we isolated 24 multidrug-resistant *Escherichia coli* genomes in Germany from humans, dogs and horses [43] (**Supplementary Table S2**). Isolates were sequenced on the Illumina MiSeq platform using the Nextera XT sequencing kit (2×250 or 2×300nt) as well as the Oxford Nanopore GridION platform using a SpotON Mk I R9 Version flow Cell (FLO-MIN106), native barcoding kit (EXP-NBD103) and 1D chemistry (SQK-LSK108). Oxford Nanopore raw data (fast5) were basecalled using Albacore (1.11.8) (https://community.nanoporetech.com). For each isolate two assemblies were performed: (i) a hybrid assembly using Unicycler v0.4.6 [44] and (ii) a short read-only assembly with SPAdes. For 21 isolates, the hybrid assembly resulted in circular chromosomes which were used as the benchmarking ground truth as the majority of remaining contigs thus originate from unclosed plasmids. The remaining 3 isolates with unclosed chromosomes were excluded from the benchmark data set, as the former requirement was not fulfilled. Short read contigs shorter than 1 kbp were discarded. The remaining contigs (n=1,337) were then aligned against closed hybrid assemblies as described above. Raw sequencing data of these 21 isolates is available as NCBI BioProjects (PRJNA505407, PRJNA387731).

## Results and Discussion

### Creation of the MPS database and RDS-based inference of contig origins

The proposed new metric RDS exploits the natural distribution biases of protein coding genes between chromosomes and plasmids to classify the origin of contigs from short-read assemblies. In order to investigate and test this rationale, we aligned a broad range of bacterial protein sequences (n=69,803,841) from UniProt’s UniRef90 protein cluster representative sequences against a set of known chromosome and plasmid reference replicons from the NCBI RefSeq and NCBI Genomes databases. 12,795,544 of these protein sequences could be aligned to at least one replicon. For each of these protein sequences a two-sided Fisher’s exact test was conducted and sequences with a p value of 1 were excluded. The remaining protein sequences (n=4,108,727) along with their RDS values, product description and sequence lengths were then used to compile the final MPS database. For 99.5% of these protein sequences (n=4,089,068) a transformed hit count ratio smaller than −0.5 (n=3,600,927) or larger than 0.5 (n=488,141) was computed, indicating a rather unequal distribution between chromosomes and plasmids (**Fig. 2**). However, only a minor fraction of 7.8% (n=322,151) of all MPS had a normalized alignment hit count sum regarding both replicon types larger than 0.001. Hence, the majority of MPS database sequences was relatively rarely detected on average. These findings endorse the incorporation of the statistical significance of each MPS replicon distribution as well as the scaling by the absolute difference of replicon hit count frequencies in order to raise the contribution of abundant protein sequences and decrease the contribution of rare protein sequences for which insufficient data is available in the reference replicon sets.

**Fig. 2.**
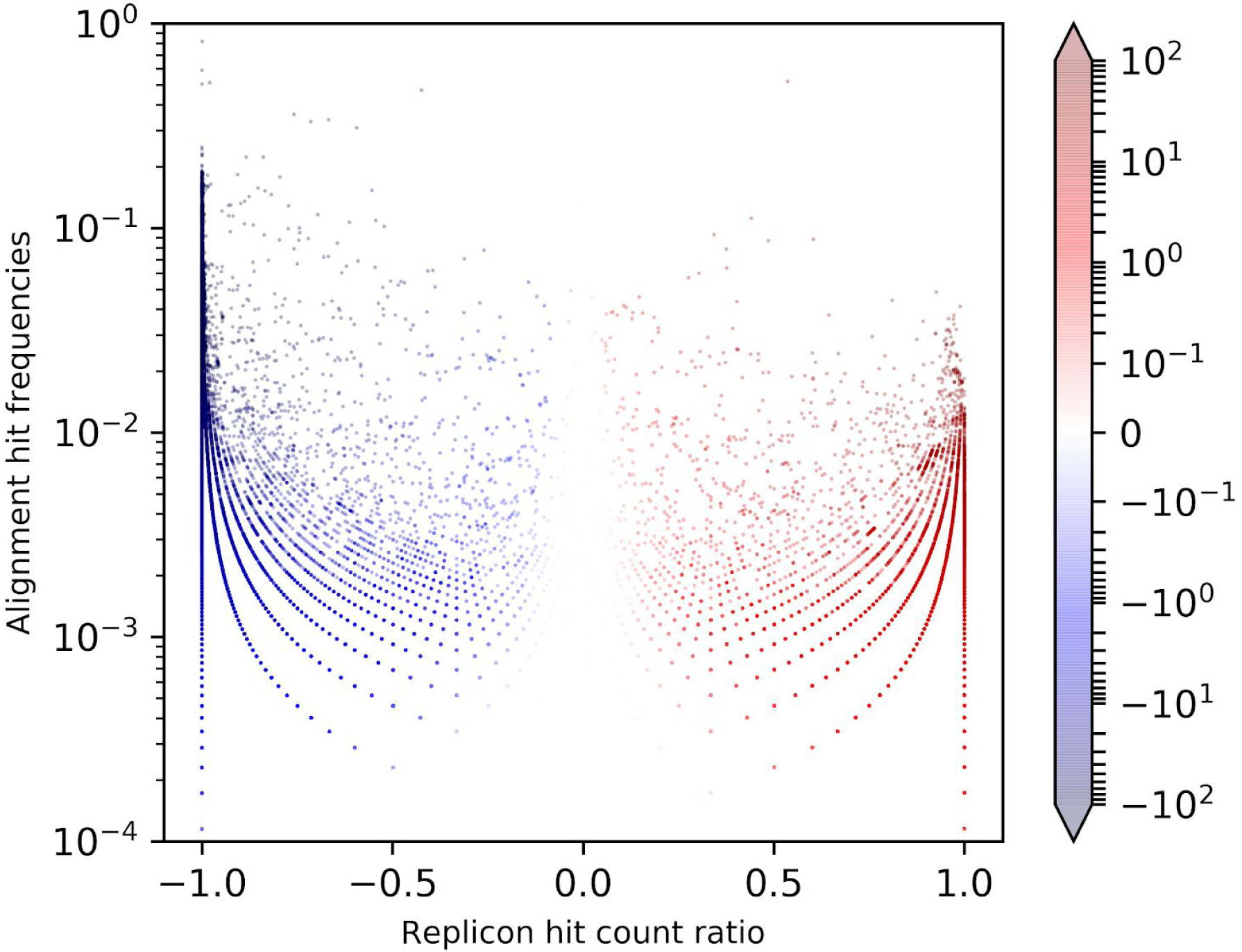
Replicon distribution and alignment hit frequencies of marker protein sequences. Shown here are summed plasmid and chromosome alignment hit frequencies per marker protein sequence plotted against plasmid/chromosome hit count ratios scaled to [-1, 1]; Hue: normalized replicon distribution score values (min=-100, max=100), hit count outliers below 10^−4^ and above 1 are discarded for the sake of readability.

In order to assess the discriminative performance of RDS regarding the replicon origin of contigs, we tested a broad range of thresholds computing sensitivity, specificity and accuracy metrics. Sensitivity, specificity and accuracy values for a range of RDS thresholds are plotted in **Fig. 3**. The sensitivity and specificity curves follow a sharp inflection point near the default RDS value, *i*.*e*. 0. We attribute this behavior to contigs harboring protein sequences which are not covered by the MPS database. To overcome this limitation and achieve both sensitive and specific classifications, we defined three distinct thresholds: (i) an SNT; (ii) an SPT; (iii) a CT set to 95% sensitivity, 99.9% specificity and the highest accuracy, respectively. Thus, contigs with an RDS smaller than the SNT can be classified as chromosomal while still retaining 95% of all plasmid contigs. Correspondingly, contigs with an RDS larger than the SPT can be classified as plasmid fragments achieving a specificity equal to or larger than 99.9%. To compute actual values for these thresholds, we conducted classifications of Monte Carlo replicon fragment simulations (n=1,564,639) by which the following values were established: SNT=-7.7, SPT=0.4 and CT=0.1 at a maximal accuracy of 84.1%. These values surround the inflection point near 0 and were henceforth used as the final discrimination thresholds in the Platon implementation.

**Fig. 3.**
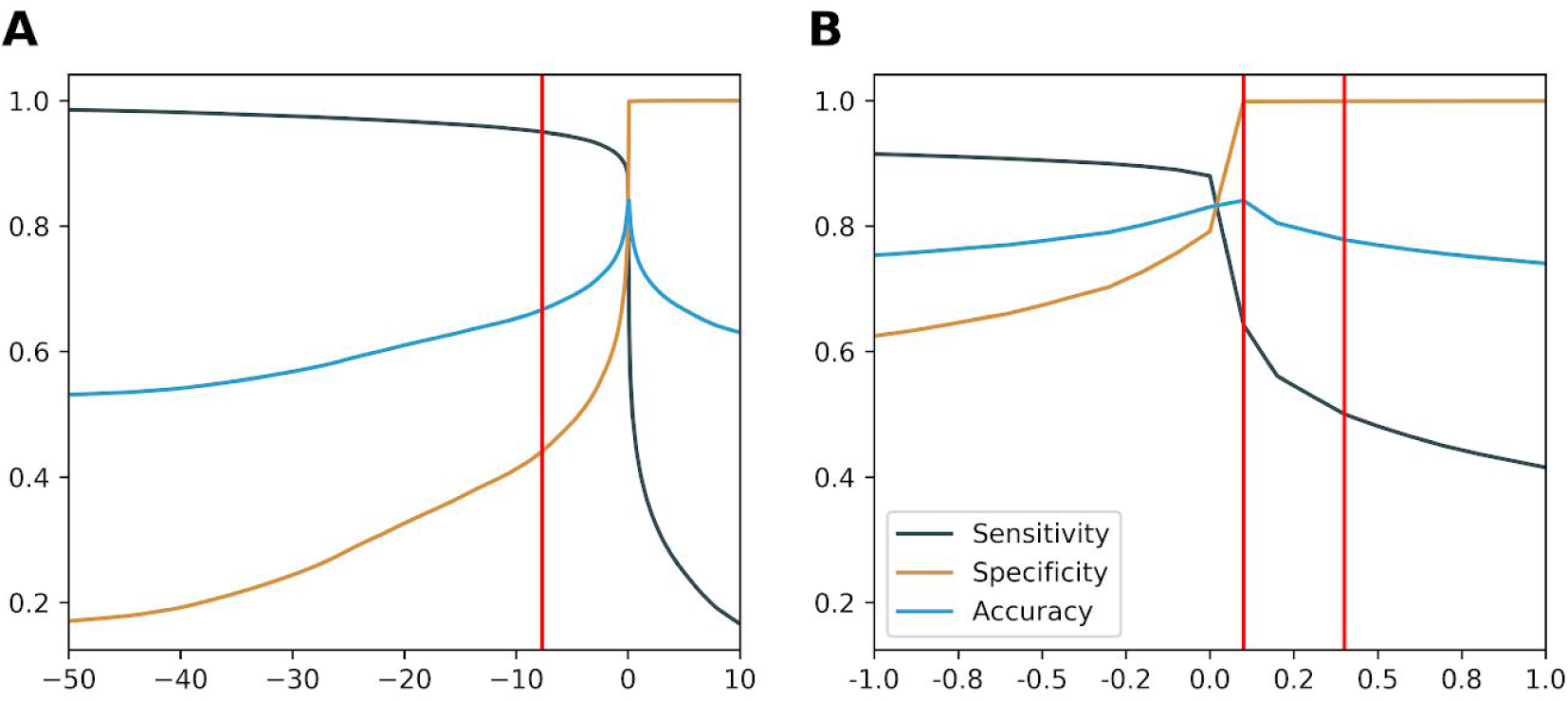
Evaluation statistics of replicon distribution score thresholds. Sensitivity, specificity and accuracy values are plotted against replicon distribution score threshold ranges; (A): overview threshold range [-50,10]; (B): detailed threshold range [-1, 1]; sensitivity in black, specificity in brown and accuracy in blue; red vertical lines from left to right: sensitivity threshold (−7.7), conservative threshold (0.1) and specificity threshold (0.4).

To finally assess the RDS based contig classification, a comprehensive performance benchmark was conducted. To do this, we created simulated short reads based on all complete NCBI RefSeq genomes (n=13,930) covering a broad range of bacterial taxa. Resulting short reads were then reassembled into contigs (n=820,392) which were aligned back to the original genomes thus creating our ground truth. This benchmark dataset comprised a total of 63,107 true plasmid contigs. All contigs were classified by their mean RDS value applying the computed SNT and SPT thresholds. This RDS workflow correctly classified 38,197 plasmid contigs and 754,082 chromosomal contigs, thus achieving an accuracy of 0.966 and a sensitivity of 0.605 as well as an F1 score of 0.731 and an MCC of 0.732 calculated upon the following confusion matrix: tp=38,197, tn=754,082, fp=3,203, fn=24,910.

Although the RDS approach achieved an accuracy of 0.966, it still misclassified 24,910 true plasmid contigs and 3,203 true chromosomal contigs. It is common knowledge that certain proteins are encoded on both replicon types as for instance relaxases and type4-coupling proteins (**T4CP**) - key proteins of integrative conjugative elements [45]. To assess the discriminative power of the RDS metric on these widespread protein classes we extracted a set of 4,683 relaxase and 2,151 T4CP clusters from the MPS database by MOBscan [39] and TXSscan [46] HMM profile searches and investigated the range of related RDS values (**Fig. S1**). 73% and 66% from the relaxase (n=3,321) and T4CP (n=1,436) protein clusters had an RDS between −0.5 and 0.5 and thus, can be considered as rather equally distributed. Small contigs solely or mainly encoding these protein sequences could therefore be especially hard to classify by the RDS metric. However, we also found 817 and 411 protein clusters which were rather chromosomally biased with a related RDS below −0.5 with extremes reaching values of −64.96 and −37.47 for the relaxases and T4CP, respectively. In addition, 445 and 304 protein clusters were rather plasmid biased with a related RDS above 0.5 with extremes reaching values of even 109.60 and 79.76 for the relaxases and T4CP, respectively. The latter protein clusters constitute about a fourth and a third of all relaxase and T4CP MPS subsets and have highly discriminative related RDS values. Hence, though there are protein classes harboring many fairly equally distributed protein clusters, *e*.*g*. the analyzed relaxases and T4CP which are often encoded in very hard to classify integrative conjugative elements, we still found MPS with a strong predictive power regarding the replicon origin of a contig.

### Performance of the entire Platon workflow

As shown in the simulated short-read benchmark the RDS metric achieved a high accuracy (ac=0.966) but rather moderate sensitivity (sn=0.605) due to the high number of false-negatives (fn=24,910). In order to increase the detection rate of true plasmid contigs, the Platon workflow additionally comprises higher-level plasmid related contig characterizations which serve as a basis for several heuristics. As both the protein homology search and the contig characterizations of large plasmids are computationally expensive, contigs larger than 500 kbp are automatically assigned to the chromosome. To assess the potentially negative impact of this heuristic on the classification performance, contig length distributions for both replicon types within the simulated short-read dataset (**Fig. S2**) were investigated. In line with the smaller plasmid contig length on average, only 119 of 63,107 plasmid contigs were actually larger than 500 kbp compared to 15,750 of 757,285 chromosome contigs. Hence, only 0.19% of all plasmid contigs were erroneously assigned to the chromosome but 99.25% of all contigs larger than 500 kbp were correctly classified by this heuristic, which thus qualifies as an eligible tradeoff between sensitivity and runtime.

To measure and assess the overall classification performance of the entire implemented workflow (**Fig. 1**), we conducted two benchmarks against contemporary command line tools: an untargeted benchmark against PlasFlow on the aforementioned simulated short-read dataset as well as a targeted benchmark against PlaScope and PlasmidFinder on sequenced *Escherichia coli* isolates.

#### Performance benchmark on taxonomically diverse simulated short-read assemblies

To assess the performance of the extended Platon workflow in an untargeted, *i*.*e*. taxon-independent setup, we conducted a comprehensive benchmark against PlasFlow, a contemporary plasmid prediction tool for metagenomics that was presented to be also eligible for the recruitment of plasmid contigs from isolates. For this benchmark, all complete bacterial NCBI RefSeq genomes (n=13,930) covering a broad range of bacterial taxa were used to simulate short reads which were *de-novo* assembled. Resulting contigs were then aligned back onto original genomes. A confusion matrix as well as common classifier performance metrics aggregated for all contigs (n=820,392) are shown in Table 1. In this benchmark Platon recruited 48,333 and PlasFlow 45,999 true plasmid contigs resulting in comparable sensitivity and negative predictive values (NPV) of 0.762 and 0.729 and 0.98 and 0.975, respectively. However, PlasFlow predicted 17 times more false positives (fp=88,712) than Platon (fp=5,277). Due to the notably lower number of false positives, Platon clearly outperformed PlasFlow in terms of accuracy, specificity, positive predictive value (PPV) as well as the balanced metrics F1 score and Matthew’s correlation coefficient (MCC). An overview of how many contigs could be classified by which RDS threshold and heuristic filter is given in Table S3.

**Table 1.** Performance benchmark results computed contig-wise on simulated short-read data.

| Metric | PlasFlow | Platon |
| --- | --- | --- |
| Accuracy | 0.871 | **0.976** |
| Sensitivity | 0.729 | **0.766** |
| Specificity | 0.883 | **0.993** |
| PPV | 0.341 | **0.902** |
| NPV | 0.975 | **0.981** |
| F1 | 0.465 | **0.828** |
| MCC | 0.440 | **0.818** |
| TP | 45,999 | **48,333** |
| TN | 668,573 | **752,080** |
| FP | 88,712 | **5,277** |
| FN | 17,108 | **14,774** |

Due to different contig lengths, the mere number of correctly classified contigs might not always be congruent with the recruited plasmid content, which could play a vital role in downstream analyses, *e*.*g*. the recruitment of plasmid-borne genes or sequence motifs, as for instance oriT and oriV. Hence, benchmarks only measuring the number of classified contigs might, to some extent, be misleading wherefore we complemented the former benchmark with a genomic content-based view calculating an additional confusion matrix based on classified DNA nucleotides (**Table S4**). **Figure 4** provides a combined view on both benchmark setups. In this complementary benchmark specificity values for PlasFlow increased from 0.883 contig-wise to 0.979 nucleotide-wise compared to stable and higher values for Platon (contig-wise=0.993; nucleotide-wise=0.995). Accuracy values also increased from 0.871 contig-wise to 0.974 nucleotide-wise for PlasFlow whereas accuracy values achieved by Platon only slightly improved (contig-wise=0.976; nucleotide-wise=0.99). Taking into account the genomic content of classified contigs revealed a performance improvement of PlasFlow in terms of accuracy and specificity but it still fell slightly behind Platon. However, PlasFlow predicted 4.3 times more false positive plasmid nucleotides (fp=1,115.3 mbp) than Platon (fp=260.9 mbp) in line with the contig-wise benchmark.

**Fig. 4.**
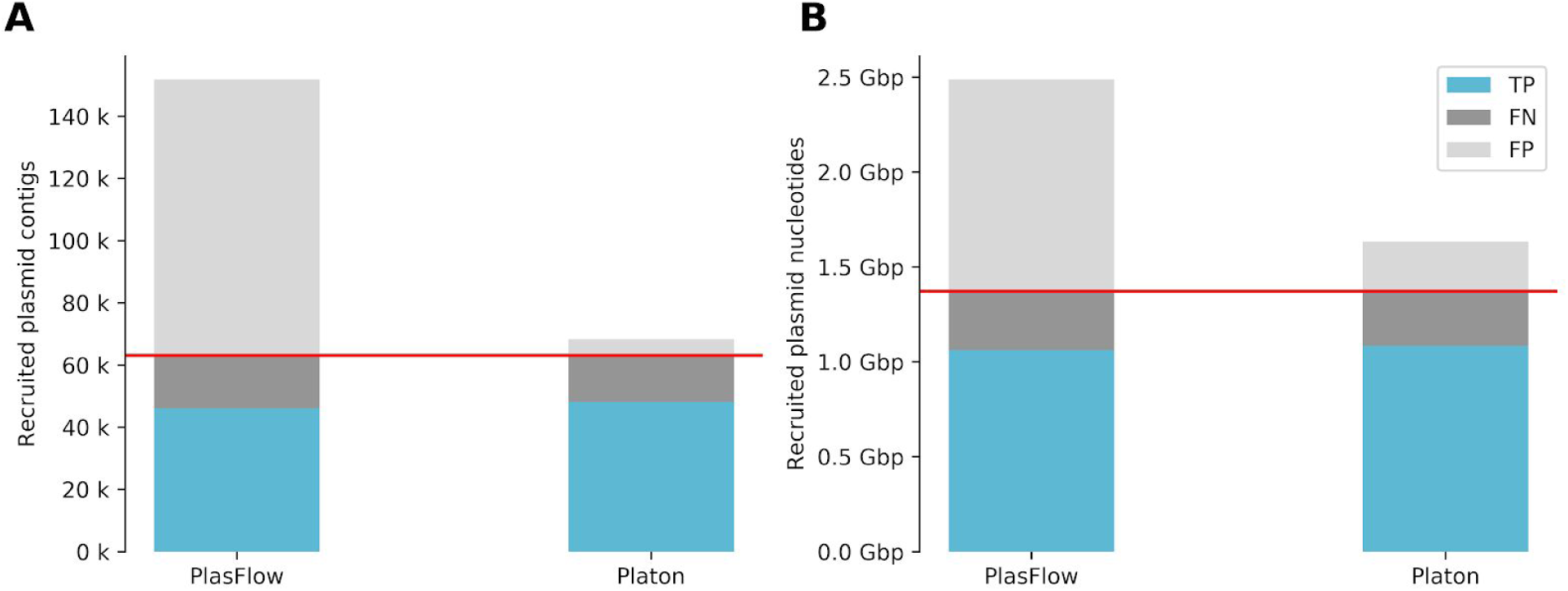
Performance benchmark metrics on simulated short read data. A performance benchmark was conducted on all complete bacterial genomes of the NCBI RefSeq database assembling simulated short reads and subsequently realigning them onto original genomes. For scaling reasons and the sake of readability, true negatives were discarded. (A) Benchmark results calculated contig-wise; Horizontal red line: total number of true plasmid contigs. (B) Benchmark results calculated nucleotide-wise; Horizontal red line: total number of true plasmid DNA nucleotides.

The taxonomic compositions of training datasets for machine-learning approaches and pre-built databases can have a severe impact on benchmark performances and the results of conducted analyses. To assess a potential bias towards certain taxa we additionally analyzed the taxonomic distribution of the recruited plasmid contigs of the simulated short-read dataset binned to the genus level (**Fig. 5**). The underlying benchmark dataset contained true plasmid contigs from 469 distinct genera and 1,234 species. From these, Platon recruited plasmid contigs from 434 genera whereas PlasFlow recruited plasmid contigs from 384 genera (**Table S5**). For both tools, the three taxa *Escherichia, Klebsiella* and *Enterococcus* accounted for nearly 40 % of the recruited sequences likewise the taxonomic profile of the underlying benchmark dataset in which aforementioned taxa accounted for 26%. On a species level, Platon and PlasFlow recruited plasmid contigs from 1,128 and 1,014 distinct species, respectively in line with aforementioned genus-level results. Although PlasFlow was developed as an untargeted tool for metagenomics, Platon recruited plasmid contigs from a wider taxonomic range, thus demonstrating the competitive edge of the taxon-independent RDS approach complemented by contig characterization heuristics.

**Fig. 5.**
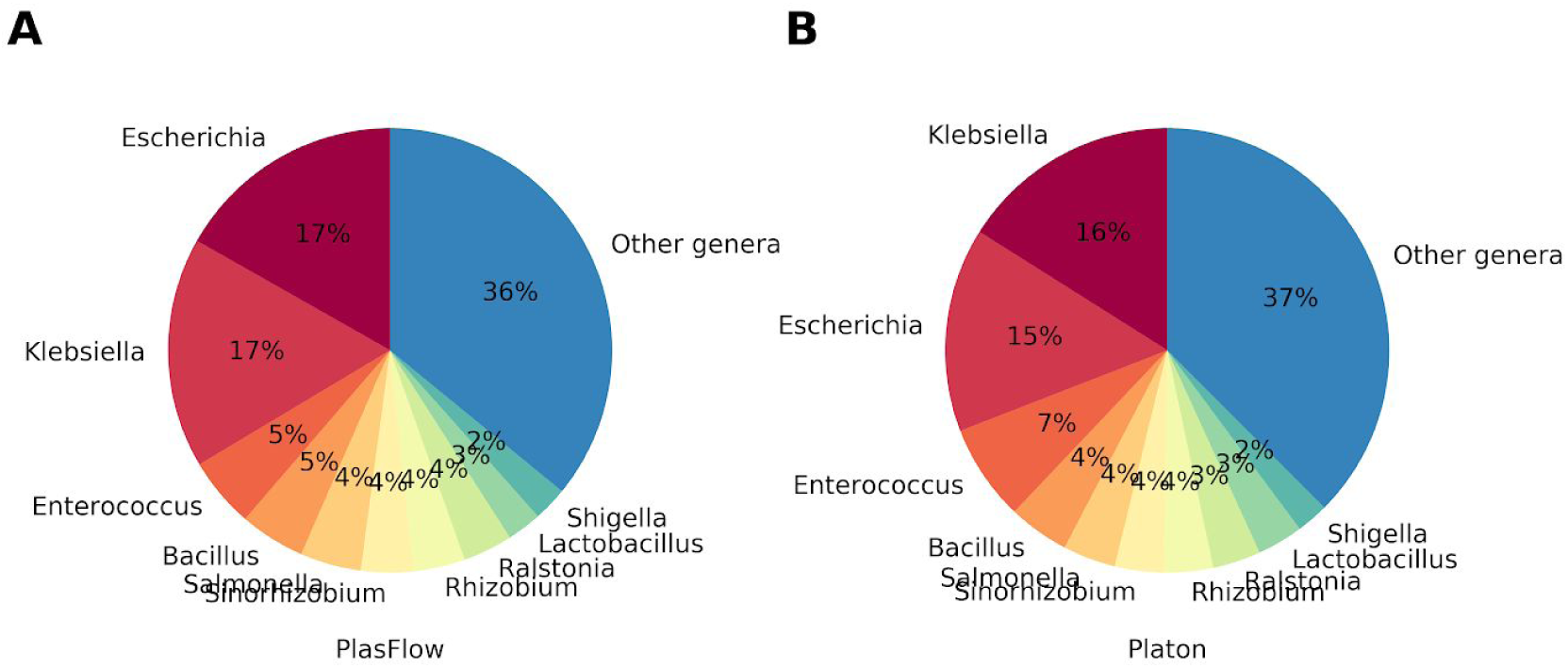
Taxonomic distribution of recruited plasmid contigs. The taxonomic distribution of recruited plasmid contigs for the simulated benchmark dataset are shown binned to the genus level. Taxa accounting for less than 2 % are grouped as “others”. (A) PlasFlow; (B) Platon.

#### Targeted performance benchmark on sequenced *Escherichia coli* isolates

Simulated data seldom reflect the existing biological and technical complexity and the plethora of potential pitfalls. Hence, we additionally benchmarked the Platon workflow on real data in a targeted setup. We compared the performance of Platon against PlaScope and PlasmidFinder which were both published as targeted approaches for the plasmid prediction within whole-genome sequencing data. PlaScope provides a pre-compiled *E. coli* database for download which was used in this benchmark and PlasmidFinder was specifically designed for the analysis of *Enterobacteriaceae* genomes. As the PlasmidFinder database is part of Platon’s contig characterization we assessed its performance to transparently compare both tools side by side. For this benchmark the genomes of 24 *E. coli* isolates were sequenced using both Illumina short-read and Oxford Nanopore long-read technologies. For 21 isolates the hybrid assemblies resulted in closed chromosomes which were used as the ground truth data. Contigs from short read-only assemblies (n=1,337) were aligned to the closed assemblies and used as the actual benchmark data. Table 2 shows the confusion matrix as well as computed benchmark metrics. PlasmidFinder achieved the lowest false positive rate (fp=14) resulting in the highest specificity of 0.987 closely followed by Platon (sp=0.966) and PlaScope (sp=0.952) but showed the lowest true positive rate (tp=57) and sensitivity (sn=0.223), thus performing worse than Platon (sn=0.699) and PlaScope (sn=0.684). Regarding accuracy, PPV, NPV, F1 score and MCC metrics Platon and PlaScope performed nearly on par, though Platon was slightly ahead on each. Both tools performed better than PlasmidFinder on these metrics. This was especially true for the balanced metrics F1 score and MCC for which Platon and Plascope clearly outperformed PlasmidFinder.

**Table 2.** Performance benchmark results contig-wise on sequenced isolate short read data.

| Metric | PlaScope | PlasmidFinder | Platon |
| --- | --- | --- | --- |
| Accuracy | 0.901 | 0.841 | **0.915** |
| Sensitivity | 0.684 | 0.223 | **0.699** |
| Specificity | 0.952 | **0.987** | 0.966 |
| PPV | 0.771 | 0.803 | **0.829** |
| NPV | 0.927 | 0.843 | **0.931** |
| F1 | 0.725 | 0.349 | **0.758** |
| MCC | 0.666 | 0.368 | **0.711** |
| TP | 175 | 57 | **179** |
| TN | 1,029 | **1,067** | 1,044 |
| FP | 52 | **14** | 37 |
| FN | 81 | 199 | **77** |

Analogically with the simulated short-read benchmark we also compared the performances of Platon, PlaScope and PlasmidFinder taking into account the amount of genomic content (Fig. 6) computed on a nucleotide-wise confusion matrix (**Table S6**). Nucleotide-wise results were in line with those calculated contig-wise: PlasmidFinder had the lowest number of false positives but also detected remarkably fewer plasmid nucleotides than PlaScope and Platon. The latter two detected a nearly equal amount of plasmid content with Platon predicting notably fewer false positives than PlaScope.

**Fig. 6.**
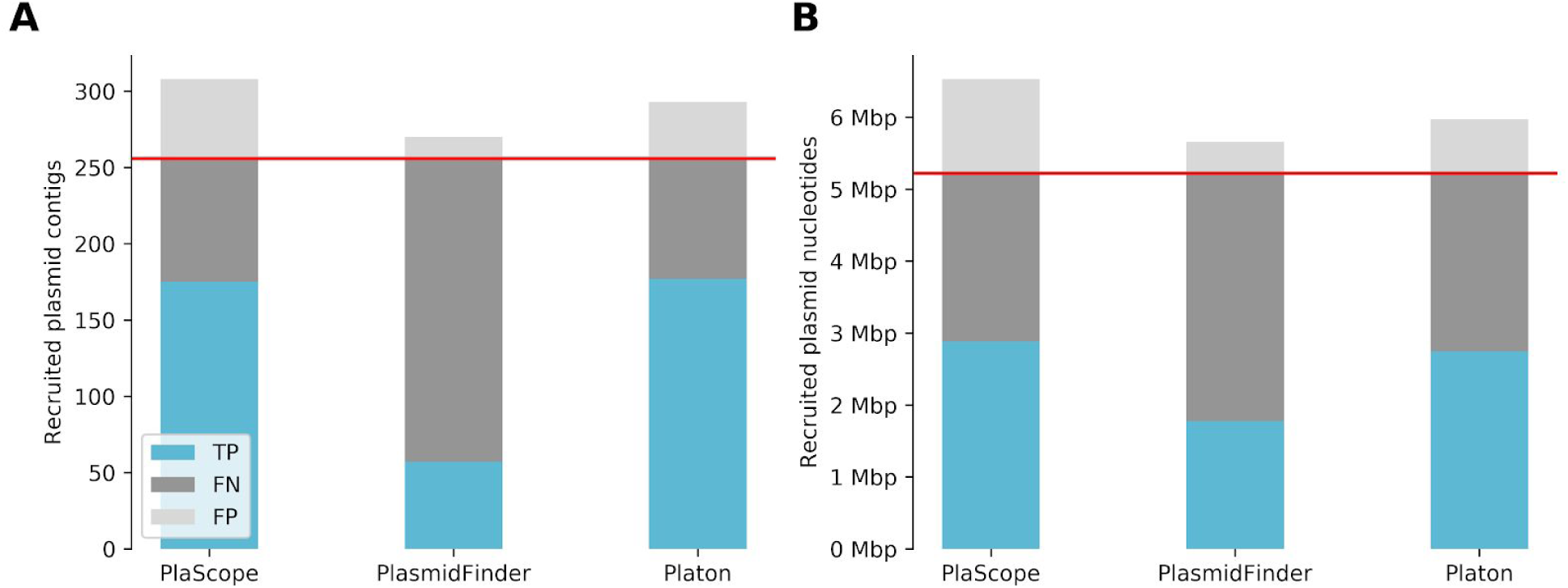
Performance benchmark metrics on real short read data. A performance benchmark was conducted on 21 *Escherichia coli* genomes for which both short read draft assemblies and complete genomes via hybrid assemblies were available. For scaling reasons and the sake of readability, true negatives were discarded. (A) Benchmark results calculated contig-wise; Horizontal red line: total number of true plasmid contigs. (B) Benchmark results calculated nucleotide-wise; Horizontal red line: total number of true plasmid DNA nucleotides.

## Conclusion

Due to the complex nature of plasmid fragments, the replicon type classification, *i*.*e*. prediction of origin, for contigs resulting from short-read draft assemblies is a difficult task. Many different methods and tools have recently been published, though few work on draft assemblies only, are implemented in a high-throughput savvy manner or provide statistically balanced predictions in an untargeted, *i*.*e*. taxon-independent manner.

To tackle this issue, we investigated the natural distribution biases of protein-coding genes between chromosomes and plasmids for a large set of protein sequences in bacteria. In this study, we defined, computed and tested statistical discrimination thresholds for the introduced new metric RDS and showed that it is a feasible approach to the problem. However, small contigs without sufficient protein sequences or contigs encoding for protein sequences which were either not covered by the MPS database or equally distributed between chromosomes and plasmids remained hard to classify correctly. However, even for the protein classes relaxases and T4CP, which are often found on notoriously hard to classify integrative conjugative elements, we found protein sequences with a strong predictive power. To mitigate these drawbacks and to improve the overall sensitivity, we complemented this approach with several heuristics exploiting higher-level plasmid-related sequence characterizations. We implemented this new workflow in a software tool called Platon and conducted benchmarks against three contemporary software tools, *i*.*e*. PlaScope, PlasFlow and PlasmidFinder on both simulated short-read data and sequenced isolates.

Analyzing a large set of diverse bacterial species Platon achieved equal sensitivity but higher accuracy and specificity than PlasFlow while the predictions made by Platon were more balanced in terms of F1 score and MCC due to a low number of false-positives.

Even though the underlying MPS database follows an untargeted approach, *i*.*e*. it is not restricted to or focused on certain taxa, Platon achieved competitive results compared to the targeted tools PlaScope and PlasmidFinder in a benchmark using real sequencing data of *E. coli* isolates. In both benchmarks Platon achieved the highest sensitivity and accuracy, thus endorsing the exploitation of the natural replicon distribution biases of protein-coding genes as an eligible method for the large-scale, high-throughput, taxon-independent prediction of plasmid-borne contigs from short-read draft assemblies.

Implemented as a multithreaded, locally-executable Linux command line application in Python 3 we also envision it as an appropriate fit for the integration into larger analysis pipelines as well as an upfront tool for subsequent plasmid specific analyses. For the sake of a streamlined integration and installation, all necessary third party executables are bundled with the software. All source code and documentation is freely available under a GPL3 license and hosted at GitHub (https://github.com/oschwengers/platon) and platon.computational.bio. For further convenience, Platon is also available as a BioConda package (`platon`) and via PyPI (`cb-platon`). A prebuilt database is hosted at Zenodo (DOI: 10.5281/zenodo.3349652).

Future developments will include the addition of new higher-level contig characterizations as well as further enhancements of applied heuristics.

## Data bibliography

1. Platon was developed as a Python 3 command line application for Linux.
2. The complete source code and documentation is available on GitHub under a GPL3 license: https://github.com/oschwengers/platon and platon.computational.bio.
3. All database versions are hosted at Zenodo: DOI 10.5281/zenodo.3349651.
4. Platon is available via bioconda package `platon`
5. Platon is available via PyPI package `cb-platon`
6. Bacterial representative sequences for UniProt’s UniRef90 protein clusters, complete bacterial genome sequences from the NCBI RefSeq database, complete plasmid sequences from the NCBI genomes plasmid section, created artificial contigs, RDS threshold metrics and raw protein replicon hit counts used to create and evaluate the marker protein sequence database are hosted at Zenodo: DOI 10.5281/zenodo.3759169s
7. 24 *Escherichia coli* isolates sequenced with short read (Illumina MiSeq) and long read sequencing technologies (Oxford Nanopore Technology GridION platform) used for real data benchmarks are available under the following NCBI BioProjects: PRJNA505407, PRJNA387731

## Funding information

This work was supported by the German Center of Infection Research (DZIF) (DZIF grant 8000 701–3 [HZI], TI06.001, 8032808811, and 8032808820 to TC); the German Network for Bioinformatics Infrastructure (de.NBI) (BMBF grant FKZ 031A533B to AG); and the German Research Foundation (DFG) (SFB-TR84 project A04 [TRR84/3 2018] to TC, KFO309 Z1 [GO 2037/5-1] to AG, SFB-TR84 project B08 [TRR84/3 2018] to TH, SFB1021 Z02 [SFB 1021/2 2017] to TH, KFO309 Z1 [HA 5225/1-1] to TH).

## Acknowledgements

The authors thank Jane Falgenhauer for her valuable ideas, testing, bug reports and fruitful feedback. Furthermore, the authors thank Christina Gerstmann for excellent technical assistance.

## Conflicts of interest

The authors declare that there are no conflicts of interest.

## Notes

http://platon.computational.bio

### Competing Interest Statement

The authors have declared no competing interest.

### Summary of Updates

This manuscript has been updated regarding new benchmark results achieved by a new software release (v1.3.1).

