## Supplementals for "Platon: identification and characterization of bacterial plasmid contigs in short-read draft assemblies exploiting protein-sequence-based replicon distribution scores"

**Supplementary Table S1.** Regular expressions used for PCLA cluster extraction for subsequent HMM creation

| Type | Regular Expression |
|---|---|
| conjugation | Tra[^ICG]<br>Trb[A-Z]<br>Trw[^ABC]<br>VirB[0-9] |
| replication | RepH<br>SopA<br>KorB<br>ParM<br>ParR<br>.*(plasmid).+(partition).*<br>.*(plasmid).+(rep) |

**Supplementary Table S2.** Isolated and sequenced *Escherichia coli* genomes used in the real data benchmark.

| Isolate | SRA Accession ID | Host | Assembled Contigs from Short Reads >= 1 kbp | Chromosome Closed in Hybrid Assemblies | Closed Plasmids in Hybrid Assemblies |
|---|---|---|---|---|---|
| H69 | SRX5007771<br>SRX5007759 | Homo sapiens | 69 | yes | 2 |
| H100 | SRX5007774<br>SRX5007760 | Homo sapiens | 44 | yes | 3 |
| H105 | SRX5002893<br>SRX5002892 | Homo sapiens | 59 | yes | 4 |
| H108 | SRX5007773<br>SRX5007761 | Homo sapiens | 69 | yes | 3 |
| H113 | SRX5007776<br>SRX5007762 | Homo sapiens | 35 | yes | 2 |
| H136 | SRX5007775<br>SRX5007763 | Homo sapiens | 77 | yes | 5 |
| H157 | SRX5007770<br>SRX5007756 | Homo sapiens | 87 | yes | 4 |
| H162 | SRX5007769<br>SRX5007757 | Homo sapiens | 56 | yes | 1 |
| H176 | SRX5007772<br>SRX5007758 | Homo sapiens | 92 | yes | 2 |
| V1 | SRX5007768<br>SRX5007764 | Canis lupus | 61 | yes | 6 |
| V8 | SRX5007790<br>SRX5007782 | Equus caballus | 46 | yes | 2 |
| V9 | SRX5007786<br>SRX5007784 | Equus caballus | 118 | yes | 5 |
| V41 | SRX5007794<br>SRX5007777 | Canis lupus | 82 | yes | 3 |
| V64 | SRX5007789<br>SRX5007780 | Canis lupus | 52 | yes | 2 |
| V71 | SRX5007788<br>SRX5007766 | Canis lupus | 90 | yes | 7 |
| V73 | SRX5007791<br>SRX5007783 | Equus caballus | 65 | yes | 2 |
| V79 | SRX6897800 | Equus caballus | 55 | no | 5 |

| | | | | | |
|---|---|---|---|---|---|
| | SRX6897801 | | | | |
| V80 | SRX5007787<br>SRX5007785 | Equus caballus | 68 | yes | 0 |
| V173 | SRX5007767<br>SRX5007765 | Equus caballus | 75 | yes | 1 |
| V177 | SRX5007793<br>SRX5007781 | Canis lupus | 66 | yes | 3 |
| V195 | SRX5007792<br>SRX5007779 | Canis lupus | 51 | yes | 4 |
| V292 | SRX5007795<br>SRX5007778 | Canis lupus | 99 | yes | 3 |
| V215 | SRX6897802<br>SRX6897803 | Equus caballus | | no | 6 |
| V232 | SRX6897804<br>SRX6897805 | Canis lupus | | no | 1 |

**Supplementary Table S3.** Number of classified contigs for each RDS and length threshold and characterization heuristic implemented in the Platon workflow for both simulated and real benchmarks.

| RDS threshold / heuristic | Simulated data | Real data |
|---|---|---|
| Length < 1kb | 0 | 0 |
| Length >= 500 kb | 15,869 | 22 |
| RDS SNT | 443,159 | 985 |
| RDS SPT | 42,669 | 241 |
| RDS CT | 50,525 | 212 |
| Circularity | 53,611 | 266 |
| Incompatibility group | 5,749 | 82 |
| Replication gene | 6,772 | 48 |
| Mobilization gene | 287 | 0 |
| OriT | 1,614 | 40 |

**Supplementary Table S4.** Confusion matrix for the untargeted simulated short-read data benchmark computed by classified genomic content measured in contig nucleotides.

| Metric | PlasFlow | Platon |
|---|---:|---:|
| TP | 1,061,149,767 | 1,087,412,371 |
| TN | 51,894,599,885 | 52,749,014,593 |
| FP | 1,115,299,457 | 260,884,749 |
| FN | 310,703,527 | 284,440,923 |

**Supplementary Table S5**. Taxa of bacterial genomes for which true plasmid contigs have been correctly identified by each tool in the simulated short-read benchmark binned to the *genus* taxon. Aggregated counts for each *genus* are provided in parenthesis.

| PlasFlow | Platon |
|---:|---:|
| *Escherichia* (7585) | *Klebsiella* (7651) |
| *Klebsiella* (7577) | *Escherichia* (7099) |
| *Enterococcus* (2309) | *Enterococcus* (3382) |
| *Bacillus* (2150) | *Bacillus* (2109) |
| *Salmonella* (2002) | *Salmonella* (1835) |
| *Sinorhizobium* (1713) | *Sinorhizobium* (1706) |
| *Rhizobium* (1707) | *Rhizobium* (1684) |
| *Ralstonia* (1633) | *Ralstonia* (1659) |
| *Lactobacillus* (1139) | *Lactobacillus* (1596) |
| *Shigella* (1124) | *Shigella* (1109) |
| *Enterobacter* (898) | *Enterobacter* (881) |
| *Xanthomonas* (653) | *Acinetobacter* (792) |
| *Acinetobacter* (647) | *Piscirickettsia* (750) |
| *Staphylococcus* (645) | *Acetobacter* (728) |
| *Acetobacter* (591) | *Xanthomonas* (641) |
| *Citrobacter* (570) | *Staphylococcus* (638) |
| *Pseudomonas* (534) | *Citrobacter* (571) |
| *Azospirillum* (504) | *Pseudomonas* (539) |
| *Piscirickettsia* (489) | *Borreliella* (503) |
| *Yersinia* (456) | *Yersinia* (464) |
| *Burkholderia* (433) | *Phaeobacter* (429) |
| *Borreliella* (336) | *Azospirillum* (418) |
| *Vibrio* (326) | *Borrelia* (380) |
| *Sphingobium* (319) | *Burkholderia* (363) |
| *Rhodococcus* (296) | *Vibrio* (339) |
| *Borrelia* (292) | *Sphingobium* (310) |
| *Lactococcus* (285) | *Lactococcus* (308) |
| *Phaeobacter* (283) | *Rhodococcus* (258) |
| *Microvirga* (260) | *Streptomyces* (258) |
| *Agrobacterium* (259) | *Paracoccus* (249) |
| *Paracoccus* (234) | *Agrobacterium* (242) |
| *Paraburkholderia* (232) | *Deinococcus* (216) |
| *Streptomyces* (208) | *Aeromonas* (200) |
| *Aeromonas* (183) | *Microvirga* (165) |
| *Deinococcus* (173) | *Clostridium* (163) |
| *Nostoc* (162) | *Nostoc* (153) |
| *Cupriavidus* (161) | *Mycobacterium* (145) |
| *Sphingomonas* (157) | *Cupriavidus* (145) |
| *Rhodobacter* (151) | *Pantoea* (139) |
| *Mycobacterium* (151) | *Pediococcus* (135) |
| *Pantoea* (143) | *Pseudonocardia* (128) |
| *Methylobacterium* (142) | *Campylobacter* (121) |
| *Novosphingobium* (128) | *Novosphingobium* (120) |
| *Komagataeibacter* (123) | *Thermus* (118) |
| *Bradyrhizobium* (114) | *Komagataeibacter* (118) |
| *Leclercia* (112) | *Sphingomonas* (114) |
| *Shewanella* (103) | *Rhodobacter* (113) |
| *Raoultella* (101) | *Paraburkholderia* (108) |

| | |
|---|---|
| *Pediococcus* (99) | *Raoultella* (106) |
| *Ensifer* (96) | *Moraxella* (102) |
| *Pandoraea* (91) | *Shewanella* (102) |
| *Clostridium* (89) | *Leuconostoc* (96) |
| *Pseudonocardia* (85) | *Ensifer* (96) |
| *Thermus* (82) | *Sulfitobacter* (95) |
| *Sulfitobacter* (80) | *Leclercia* (92) |
| Acaryochloris (75) | *Methylobacterium* (91) |
| *Ochrobactrum* (69) | *Bradyrhizobium* (91) |
| Campylobacter (68) | Listeria (82) |
| *Acidiphilium* (67) | *Arsenophonus* (79) |
| *Serratia* (66) | *Legionella* (77) |
| *Mesorhizobium* (62) | *Gloeothece* (77) |
| *Gloeothece* (62) | *Acaryochloris* (76) |
| *Roseomonas* (61) | *Pandoraea* (73) |
| Photobacterium (60) | *Candidatus* (71) |
| *Arsenophonus* (59) | *Sphingopyxis* (67) |
| *Methylorubrum* (57) | *Synechococcus* (65) |
| *Sphingopyxis* (57) | *Paenibacillus* (62) |
| *Kozakia* (55) | *Ochrobactrum* (60) |
| Arthrobacter (54) | *Arthrobacter* (60) |
| *Leptolyngbya* (51) | *Chlamydia* (59) |
| *Leuconostoc* (50) | *Photobacterium* (59) |
| *Enterobacteriaceae* (49) | *Serratia* (58) |
| *Shinella* (48) | *Acidiphilium* (57) |
| *Phytobacter* (48) | *Synechocystis* (57) |
| *Listeria* (47) | *Leptolyngbya* (56) |
| *Proteus* (44) | *Azotobacter* (51) |
| *Cronobacter* (43) | *Kozakia* (51) |
| Candidatus (40) | *Cronobacter* (50) |
| *Gluconobacter* (39) | *Zymomonas* (48) |
| Mycoplasma (39) | *Enterobacteriaceae* (47) |
| *Synechococcus* (38) | *Aminobacter* (46) |
| *Moraxella* (37) | *Phytobacter* (46) |
| *Synechocystis* (36) | *Proteus* (43) |
| *Haematobacter* (36) | *Methylorubrum* (42) |
| *Helicobacter* (35) | *Clavibacter* (41) |
| *Aminobacter* (33) | *Shinella* (41) |
| *Paenibacillus* (32) | *Mesorhizobium* (40) |
| *Acidovorax* (32) | *Acidovorax* (37) |
| *Corynebacterium* (31) | *Roseomonas* (37) |
| *Mycolicibacterium* (31) | *Geobacillus* (36) |
| *Streptococcus* (30) | *Corynebacterium* (36) |
| *Neorhizobium* (29) | *Gluconobacter* (35) |
| *Carnobacterium* (29) | *Neisseria* (34) |
| *Croceicoccus* (29) | *Leptospira* (33) |
| *Azotobacter* (28) | *Weissella* (31) |
| *Geobacillus* (28) | *Acidithiobacillus* (31) |
| *Erwinia* (26) | *Rickettsia* (31) |
| *Celeribacter* (25) | *Helicobacter* (30) |
| *Antarctobacter* (24) | *Haematobacter* (30) |
| *Legionella* (23) | *Psychrobacter* (29) |
| *Aliivibrio* (23) | *Meiothermus* (29) |
| *Oscillatoria* (23) | *Leisingera* (29) |
| *Chlamydia* (22) | *Celeribacter* (29) |
| *Xylella* (22) | *Planococcus* (28) |

| | |
|---|---|
| *Stanieria* (22) | *Calothrix* (28) |
| *Aureimonas* (22) | *Stanieria* (28) |
| Confluentimicrobium (22) | *Polaromonas* (28) |
| *Phyllobacterium* (22) | *Erwinia* (27) |
| *Zymomonas* (21) | *Carnobacterium* (27) |
| *Xenorhabdus* (21) | *Oscillatoria* (27) |
| *Nitrobacter* (21) | *Streptococcus* (26) |
| *Rahnella* (20) | *Pseudanabaena* (26) |
| *Rhodovulum* (20) | *Xylella* (25) |
| *Octadecabacter* (20) | *Antarctobacter* (25) |
| *Aromatoleum* (20) | *Yangia* (25) |
| *Rhizorhabdus* (20) | *Indioceanicola* (25) |
| *Neisseria* (19) | *Rhodovulum* (24) |
| *Gordonia* (19) | *Croceicoccus* (24) |
| *Yangia* (19) | *Neorhizobium* (23) |
| *Epibacterium* (19) | *Mycolicibacterium* (23) |
| *Buchnera* (18) | *Martelella* (23) |
| *Martelella* (18) | *Cyanothece* (23) |
| *Indioceanicola* (18) | *Bacteroides* (22) |
| *Rippkaea* (18) | *Ruminococcus* (22) |
| *Leptospira* (17) | *Rippkaea* (22) |
| *Weissella* (17) | *Buchnera* (21) |
| *Nocardia* (17) | *Aliivibrio* (21) |
| *Psychrobacter* (17) | *Anabaena* (21) |
| *Thioclava* (17) | *Granulicella* (20) |
| *Edwardsiella* (16) | *Confluentimicrobium* (20) |
| *Anabaena* (16) | *Pseudoalteromonas* (19) |
| *Cyanothece* (16) | *Marinovum* (19) |
| *Alteromonas* (15) | *Deferribacter* (19) |
| *Rickettsia* (15) | *Phyllobacterium* (19) |
| *Polaromonas* (15) | *Methylosinus* (18) |
| *Sagittula* (15) | *Ilyobacter* (18) |
| *Methylosinus* (14) | *Aromatoleum* (18) |
| *Planococcus* (14) | *Salipiger* (18) |
| *Marinobacter* (14) | *Aureimonas* (18) |
| *Calothrix* (14) | *Geminocystis* (18) |
| *Leisingera* (14) | *Peptoclostridium* (17) |
| *Dinoroseobacter* (14) | *Ketogulonicigenium* (17) |
| *Salipiger* (14) | *Epibacterium* (17) |
| *Pelagibaca* (14) | *Pelagibaca* (17) |
| *Defluviimonas* (14) | *Xenorhabdus* (16) |
| *Spiroplasma* (13) | *Desulfovibrio* (16) |
| *Clavibacter* (13) | *Rhizorhabdus* (16) |
| *Microcoleus* (13) | *Sedimentitalea* (16) |
| *Sedimentitalea* (13) | *Edwardsiella* (15) |
| *Metakosakonia* (13) | *Rahnella* (15) |
| *Marinovum* (12) | *Pseudarthrobacter* (15) |
| *Pseudanabaena* (12) | *Sagittula* (15) |
| *Alicycliphilus* (12) | *Pseudorhodobacter* (15) |
| *Gemmobacter* (12) | *Alteromonas* (14) |
| *Francisella* (11) | *Nocardia* (14) |
| *Acidithiobacillus* (11) | *Salinibacter* (14) |
| *Oligotropha* (11) | *Gemmatirosa* (14) |
| *Macrococcus* (11) | *Gemmobacter* (14) |
| *Kosakonia* (11) | *Metakosakonia* (14) |
| *Crinalium* (11) | *Fusobacterium* (13) |

| | |
|---|---|
| *Trichormus* (11) | *Nitrobacter* (13) |
| *Chelativorans* (11) | *Selenomonas* (13) |
| *Bosea* (11) | *Lysinibacillus* (13) |
| *Geminocystis* (11) | *Dinoroseobacter* (13) |
| *Frondihabitans* (11) | *Methylomonas* (13) |
| *Pseudoalteromonas* (10) | *Microcoleus* (13) |
| *Pseudarthrobacter* (10) | *Francisella* (12) |
| *Chondrocystis* (10) | *Coxiella* (12) |
| *Buttiauxella* (10) | *Parageobacillus* (12) |
| *Pseudorhodobacter* (10) | *Mycoplasma* (12) |
| *Lysinibacillus* (9) | *Marinobacter* (12) |
| *Ruegeria* (9) | *Octadecabacter* (12) |
| *Tistrella* (9) | *Ruegeria* (12) |
| *Yoonia* (9) | *Methylocystis* (12) |
| *Massilia* (9) | *Crinalium* (12) |
| *Hymenobacter* (9) | *Defluviimonas* (12) |
| *Niveispirillum* (9) | *Acidisarcina* (12) |
| *Acidisarcina* (9) | *Nitrosomonas* (11) |
| *Bifidobacterium* (8) | *Rubrobacter* (11) |
| *Mycobacteroides* (8) | *Asticcacaulis* (11) |
| *Meiothermus* (8) | *Chondrocystis* (11) |
| *Ketogulonicigenium* (8) | *Treponema* (10) |
| Dietzia (8) | *Chelativorans* (10) |
| *Cedecea* (8) | *Phenylobacterium* (10) |
| *Deferribacter* (8) | *Halomonas* (10) |
| *Granulicella* (8) | *Thioclava* (10) |
| *Morganella* (7) | *Citricoccus* (10) |
| *Bacteroides* (7) | *Xanthobacter* (9) |
| *Virgibacillus* (7) | *Bifidobacterium* (9) |
| *Tetragenococcus* (7) | *Achromobacter* (9) |
| *Methylibium* (7) | *Methylibium* (9) |
| *Methylocystis* (7) | *Sulfuricurvum* (9) |
| *Cryobacterium* (7) | *Alicycliphilus* (9) |
| *Blastomonas* (7) | *Yoonia* (9) |
| *Xanthobacter* (6) | *Kosakonia* (9) |
| *Pectobacterium* (6) | *Massilia* (9) |
| *Providencia* (6) | *Hymenobacter* (9) |
| *Ruminococcus* (6) | *Bosea* (9) |
| *Parageobacillus* (6) | *Niveispirillum* (9) |
| *Anoxybacillus* (6) | *Hoeflea* (9) |
| *Paenarthrobacter* (6) | *Planctomyces* (9) |
| *Rhodoferax* (6) | *Providencia* (8) |
| *Kocuria* (6) | *Clostridioides* (8) |
| *Halomonas* (6) | *Gordonia* (8) |
| *Rhizobiales* (6) | Haliscomenobacter (8) |
| *Porphyrobacter* (6) | *Roseobacter* (8) |
| *Citricoccus* (6) | *Oligotropha* (8) |
| *Coxiella* (5) | *Pannonibacter* (8) |
| *Clostridioides* (5) | *Tistrella* (8) |
| *Roseobacter* (5) | *Frondihabitans* (8) |
| *Thauera* (5) | *Runella* (8) |
| *Achromobacter* (5) | *Buttiauxella* (8) |
| *Methylocella* (5) | *Pectobacterium* (7) |
| *Aster* (5) | *Morganella* (7) |
| *Hoeflea* (5) | *Rhodothermus* (7) |
| *Microbacterium* (5) | *Melissococcus* (7) |

| | |
|---|---|
| *Rhodobacteraceae* (5) | *Myroides* (7) |
| *Acidibrevibacterium* (5) | *Chryseobacterium* (7) |
| *Tabrizicola* (5) | *Dietzia* (7) |
| *Crocosphaera* (5) | *Trichormus* (7) |
| *Beijerinckia* (4) | *Kocuria* (7) |
| *Plesiomonas* (4) | *Acidibrevibacterium* (7) |
| *Actinobacillus* (4) | *Virgibacillus* (6) |
| *Pasteurella* (4) | *Desulfobacterium* (6) |
| *Nitrosomonas* (4) | *Arcobacter* (6) |
| *Thiomonas* (4) | *Lawsonia* (6) |
| *Selenomonas* (4) | *Paenarthrobacter* (6) |
| *Allochromatium* (4) | *Tetragenococcus* (6) |
| *Microcystis* (4) | *Sodalis* (6) |
| *Brevibacillus* (4) | *Caulobacter* (6) |
| *Kitasatospora* (4) | *Macrococcus* (6) |
| *Tsukamurella* (4) | *Rhodoferax* (6) |
| *Lawsonia* (4) | *Simkania* (6) |
| *Gluconacetobacter* (4) | *Amycolatopsis* (6) |
| *Glutamicibacter* (4) | *Cedecea* (6) |
| *Nodularia* (4) | *Thalassospira* (6) |
| *Myroides* (4) | *Methylocella* (6) |
| *Asticcacaulis* (4) | *Singulisphaera* (6) |
| *Desulfovibrio* (4) | *Aquabacterium* (6) |
| *Labrenzia* (4) | *Rhodobacteraceae* (6) |
| *Phenylobacterium* (4) | *Clostridiaceae* (6) |
| *Exiguobacterium* (4) | *Lelliottia* (6) |
| *Oscillibacter* (4) | *Tabrizicola* (6) |
| *Vagococcus* (4) | *Plesiomonas* (5) |
| *Neokomagataea* (4) | *Nitrosococcus* (5) |
| *Plautia* (4) | *Nitrosospira* (5) |
| *Swingsia* (4) | *Pelobacter* (5) |
| *Hartmannibacter* (4) | *Mycobacteroides* (5) |
| *Clostridiaceae* (4) | *Butyrivibrio* (5) |
| *Simplicispira* (4) | *Chroococcidiopsis* (5) |
| *Bordetella* (3) | *Labrenzia* (5) |
| *Hafnia* (3) | *Roseovarius* (5) |
| *Fusobacterium* (3) | *Tateyamaria* (5) |
| *Rhodospirillum* (3) | *Exiguobacterium* (5) |
| *Halobacillus* (3) | *Oscillibacter* (5) |
| *Desulfobacterium* (3) | *Flammeovirga* (5) |
| *Bartonella* (3) | *Thermaerobacter* (5) |
| *Sodalis* (3) | *Porphyrobacter* (5) |
| *Pannonibacter* (3) | *Brachyspira* (4) |
| *Salinibacter* (3) | *Hydrogenophilus* (4) |
| *Roseovarius* (3) | *Thiomonas* (4) |
| *Thalassospira* (3) | *Rhodospirillum* (4) |
| *Alicyclobacillus* (3) | *Halobacillus* (4) |
| *Chelatococcus* (3) | *Prevotella* (4) |
| *Salimicrobium* (3) | *Geobacter* (4) |
| *Haematospirillum* (3) | *Glutamicibacter* (4) |
| *Erythrobacter* (3) | *Thauera* (4) |
| *Nostocales* (3) | *Curtobacterium* (4) |
| *Glaesserella* (3) | *Thioflavicoccus* (4) |
| *Lelliottia* (3) | *Bartonella* (4) |
| *Thiomicrorhabdus* (3) | *Blattabacterium* (4) |
| *Hydrocarboniclastica* (3) | *Geoalkalibacter* (4) |

| | |
|---|---|
| *Vitreoscilla* (2) | *Neokomagataea* (4) |
| *Hydrogenophilus* (2) | *Cryobacterium* (4) |
| *Sebaldella* (2) | *Azoarcus* (4) |
| *Finegoldia* (2) | *Gloeocapsa* (4) |
| *Cutibacterium* (2) | *Haematospirillum* (4) |
| *Haliscomenobacter* (2) | *Blastomonas* (4) |
| *Brochothrix* (2) | *Cnuibacter* (4) |
| *Pelobacter* (2) | *Microbacterium* (4) |
| *Caldicellulosiruptor* (2) | *Actinobacillus* (3) |
| *Desulfohalobium* (2) | *Allochromatium* (3) |
| *Prevotella* (2) | *Microcystis* (3) |
| *Chroococcidiopsis* (2) | *Brevibacillus* (3) |
| *Thioflavicoccus* (2) | *Tsukamurella* (3) |
| *Moritella* (2) | *Spiroplasma* (3) |
| *Simkania* (2) | *Caldicellulosiruptor* (3) |
| *endosymbiont* (2) | *Desulfohalobium* (3) |
| *Caulobacter* (2) | *Nodularia* (3) |
| *Rivularia* (2) | *Aster* (3) |
| *Cyanobacterium* (2) | *Thermovirga* (3) |
| *Singulisphaera* (2) | *Thermobacillus* (3) |
| *Hoyosella* (2) | *Cyanobacterium* (3) |
| *Azoarcus* (2) | *Vagococcus* (3) |
| *Polymorphum* (2) | *Salimicrobium* (3) |
| *Dickeya* (2) | *Neochlamydia* (3) |
| *Gloeocapsa* (2) | *Simplicispira* (3) |
| *Aquabacterium* (2) | *Chromobacterium* (3) |
| *Halocynthiibacter* (2) | *Thiomicrorhabdus* (3) |
| *Euzebya* (2) | *Silvanigrellales* (3) |
| *Cnuibacter* (2) | *Crocosphaera* (3) |
| *Planctomyces* (2) | *Vitreoscilla* (2) |
| *Nitratireductor* (2) | *Bordetella* (2) |
| *Brachybacterium* (2) | *Beijerinckia* (2) |
| *Gammaproteobacteria* (2) | *Hafnia* (2) |
| *Sterolibacteriaceae* (2) | *Pasteurella* (2) |
| *Thermaerobacter* (2) | *Sebaldella* (2) |
| *Comamonas* (1) | *Finegoldia* (2) |
| *Alcaligenes* (1) | *Dermacoccus* (2) |
| *Histophilus* (1) | *Streptosporangium* (2) |
| *Gallibacterium* (1) | *Gluconacetobacter* (2) |
| *Marivirga* (1) | *Streptobacillus* (2) |
| *Rhodopseudomonas* (1) | *Sinomonas* (2) |
| *Prosthecochloris* (1) | *Tatumella* (2) |
| *Nitrosospira* (1) | *Pseudodesulfovibrio* (2) |
| *Dermacoccus* (1) | *Desulfocapsa* (2) |
| *Brevibacterium* (1) | *endosymbiont* (2) |
| *Peptoclostridium* (1) | *Cardinium* (2) |
| *Acidipropionibacterium* (1) | *Methylovorus* (2) |
| *Desulfurella* (1) | *Jannaschia* (2) |
| *Zymobacter* (1) | *Anoxybacillus* (2) |
| *Sinomonas* (1) | *Advenella* (2) |
| *Rubrobacter* (1) | *Rivularia* (2) |
| *Hydrogenophaga* (1) | *Natranaerobius* (2) |
| *Wigglesworthia* (1) | *Thioalkalivibrio* (2) |
| *Tatumella* (1) | *Maritalea* (2) |
| *Waddlia* (1) | *Calditerrivibrio* (2) |
| *Mannheimia* (1) | *Rufibacter* (2) |

| | |
|---|---|
| *Solibacillus* (1) | *Opitutaceae* (2) |
| *Brachyspira* (1) | *Desulfosporosinus* (2) |
| *Desulfotalea* (1) | *Plautia* (2) |
| *Halobacteriovorax* (1) | *Halioglobus* (2) |
| *Kineococcus* (1) | *Polymorphum* (2) |
| *Rummeliibacillus* (1) | *Dickeya* (2) |
| *Cardinium* (1) | *Altererythrobacter* (2) |
| *Methylovorus* (1) | *Capnocytophaga* (2) |
| *Jannaschia* (1) | *Paludisphaera* (2) |
| *Photorhabdus* (1) | *Cetia* (2) |
| *Tateyamaria* (1) | *Hartmannibacter* (2) |
| *Advenella* (1) | *Rickettsiales* (2) |
| *Geobacter* (1) | *Erythrobacter* (2) |
| *Verminephrobacter* (1) | *Nostocales* (2) |
| *Natranaerobius* (1) | *Thalassococcus* (2) |
| *Thioalkalivibrio* (1) | *Glaesserella* (2) |
| *Tessaracoccus* (1) | *Gammaproteobacteria* (2) |
| *Nitrosococcus* (1) | *Catenovulum* (2) |
| *Prauserella* (1) | *Humibacter* (2) |
| *Allofrancisella* (1) | *Planctopirus* (1) |
| *Frankia* (1) | *Isosphaera* (1) |
| *Jeotgalibaca* (1) | *Comamonas* (1) |
| *Methylophaga* (1) | *Alcaligenes* (1) |
| *Arcobacter* (1) | *Histophilus* (1) |
| *Pusillimonas* (1) | *Gallibacterium* (1) |
| *Moorea* (1) | *Herbaspirillum* (1) |
| *Catharanthus* (1) | *Marivirga* (1) |
| *Mucilaginibacter* (1) | *Saprospira* (1) |
| *Cycloclasticus* (1) | *Prosthecochloris* (1) |
| *Paludisphaera* (1) | *Gottschalkia* (1) |
| *Geosporobacter* (1) | *Kitasatospora* (1) |
| *Sedimenticola* (1) | *Hirschia* (1) |
| *Aquitalea* (1) | *Brochothrix* (1) |
| *Psychromicrobium* (1) | *Turneriella* (1) |
| *Spongiibacter* (1) | *Desulfurella* (1) |
| *Magnetospirillum* (1) | *Zymobacter* (1) |
| *Agarilytica* (1) | *Eubacterium* (1) |
| *Fischerella* (1) | *Rothia* (1) |
| *Paraphotobacterium* (1) | *Hydrogenophaga* (1) |
| *Sphingosinicella* (1) | *Wigglesworthia* (1) |
| *Amycolatopsis* (1) | *Flavobacterium* (1) |
| *Tenericutes* (1) | *Waddlia* (1) |
| *Marivivens* (1) | *Mannheimia* (1) |
| *Sporosarcina* (1) | *Moritella* (1) |
| *Sulfuriferula* (1) | *Desulfotalea* (1) |
| *Thalassococcus* (1) | *Halobacteriovorax* (1) |
| *Ahniella* (1) | *Kineococcus* (1) |
| *Mycetocola* (1) | *Marinitoga* (1) |
| *Butyricimonas* (1) | *Carboxydocella* (1) |
| *Miniimonas* (1) | *Xylanimonas* (1) |
| *Catenovulum* (1) | *Thermovibrio* (1) |
| *Runella* (1) | *Methylomicrobium* (1) |
| *Humibacter* (1) | *Collimonas* (1) |
| *Flammeovirga* (1) | *Photorhabdus* (1) |
| *Xylanibacterium* (1) | *Persephonella* (1) |
| *Xanthomonadaceae* (1) | *Pontibacter* (1) |

| | |
|---|---|
| *Rhodopseudomonas* (1) | *Verminephrobacter* (1) |
| *Tatumella* (1) | *Alicyclobacillus* (1) |
| *Nitrosospira* (1) | *Oceanimonas* (1) |
| *Fischerella* (1) | *Prauserella* (1) |
| *Mannheimia* (1) | *Pelagibacterium* (1) |
| *Photorhabdus* (1) | *Phycisphaera* (1) |
| *Spongiibacter* (1) | *Allofrancisella* (1) |
| *Jannaschia* (1) | *Hoyosella* (1) |
| Methylovorus (1) | *Sulfuricella* (1) |
| *Thalassococcus* (1) | *Frankia* (1) |
| *Alcaligenes* (1) | *Jeotgalibaca* (1) |
| *Catenovulum* (1) | *Verrucosispora* (1) |
| *Zymobacter* (1) | *Pusillimonas* (1) |
| Tenericutes (1) | *Thiolapillus* (1) |
| *Xanthomonadaceae* (1) | *Elizabethkingia* (1) |
| *Mucilaginibacter* (1) | *Mesotoga* (1) |
| *Jeotgalibaca* (1) | *Catharanthus* (1) |
| *Mycetocola* (1) | *Magnetospira* (1) |
| *Psychromicrobium* (1) | *Swingsia* (1) |
| *Hydrogenophaga* (1) | *Mucilaginibacter* (1) |
| | *Cycloclasticus* (1) |
| | *Serpentinomonas* (1) |
| | *Erysipelothrix* (1) |
| | *Sedimenticola* (1) |
| | *Halocynthiibacter* (1) |
| | *Aquitalea* (1) |
| | *Psychromicrobium* (1) |
| | *Spongiibacter* (1) |
| | *Mitsuaria* (1) |
| | *Magnetospirillum* (1) |
| | *Chelatococcus* (1) |
| | *Agarilytica* (1) |
| | *Fischerella* (1) |
| | *Paraphotobacterium* (1) |
| | *Nitratireductor* (1) |
| | *Sphingosinicella* (1) |
| | *Acetobacteraceae* (1) |
| | *Marivivens* (1) |
| | *Sporosarcina* (1) |
| | *Sulfuriferula* (1) |
| | *Brachybacterium* (1) |
| | *Ahniella* (1) |
| | *Sphingorhabdus* (1) |
| | *Butyricimonas* (1) |
| | *Sterolibacteriaceae* (1) |
| | *Hydrocarboniclastica* (1) |
| | *Oenococcus* (1) |
| | *Thermoactinomycetaceae* (1) |
| | *Streptomonospora* (1) |
| | *Rhizobiales* (1) |

**Supplementary Table S6.** Confusion matrix for the targeted real short-read/long-read hybrid data benchmark computed by classified genomic content measured in contig nucleotides.

| Metric | PlaScope | PlasmidFinder | Platon |
|---|---|---|---|
| TP | 2,884,199 | 1,776,553 | 2,745,897 |
| TN | 97,966,253 | 98,841,671 | 98,525,184 |
| FP | 1,309,315 | 433,897 | 750,384 |
| FN | 2,337,708 | 3,445,354 | 2,476,010 |

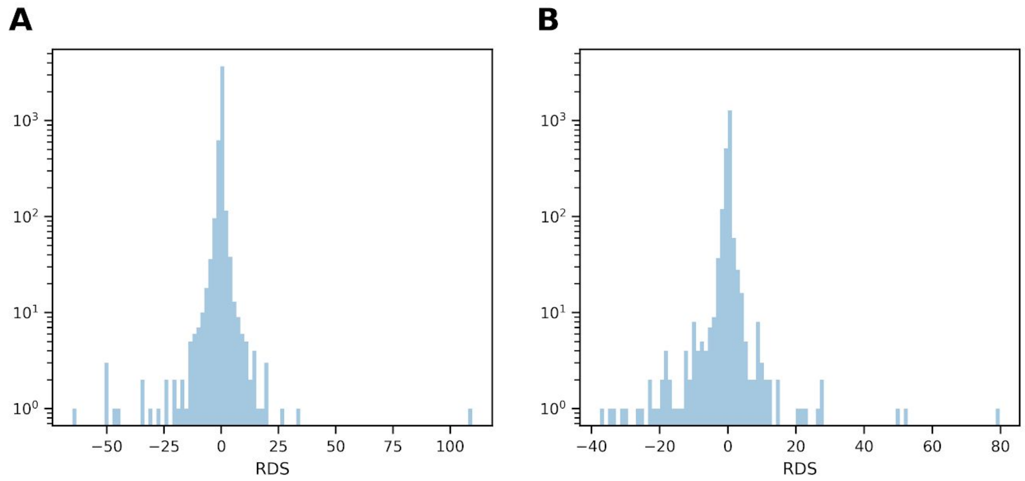


**Supplementary Figure S1**. Histogram of RDS values for (A) relaxase and (B) type 4-coupling proteins.

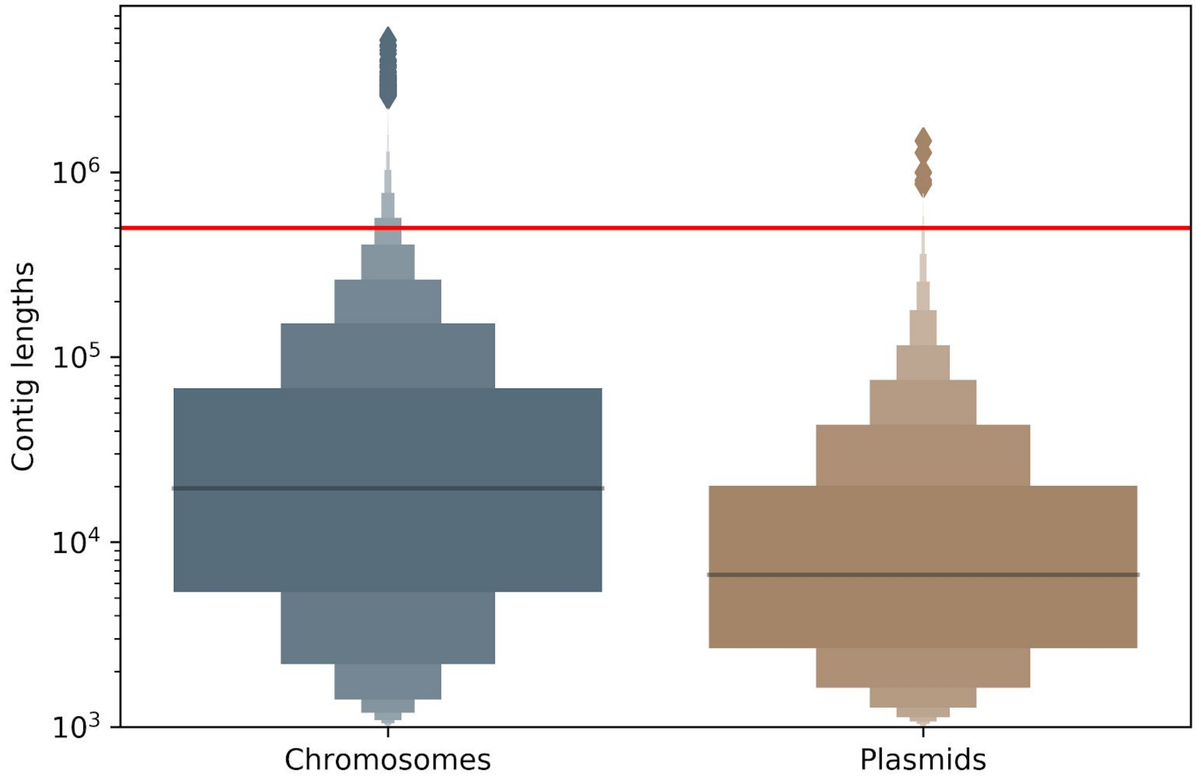


**Supplementary Figure S2**. Length distribution of chromosome and plasmid contigs resulted from simulated short-read assemblies. Outliers are shown as diamonds; horizontal red line: implemented contig length heuristic threshold (n=500,000 bp) as applied in the platon workflow.
